# Molecular Evolution of Aryl Hydrocarbon Receptor Signaling Pathway Genes

**DOI:** 10.1101/2022.07.13.499904

**Authors:** Diksha Bhalla, Vera van Noort

## Abstract

The Aryl hydrocarbon receptor is an ancient transcriptional factor originally discovered as a sensor of dioxin. In addition to its function as receptor of environmental toxins, it plays an important role in development. Although a significant amount of research has been carried out to understand AHR signal transduction pathway and its involvement in species susceptibility to environmental toxins, none of them to date has comprehensively studied its evolutionary origins. Studying evolutionary origins of molecules can inform ancestral relationships of genes. Vertebrate genome has been shaped by two rounds of whole-genome duplications at the base of vertebrate evolution approximately 600 million years ago (Mya), followed by lineage specific gene losses, which often complicate assignment of functional homology. It is crucial to understand evolutionary origins of this transcription factor and its partners, in order to distinguish orthologs from ancient non-orthologous homologs. In this study, we have investigated evolutionary origins of proteins involved in the AHR pathway. Our results provide evidence of gene loss and duplications, crucial for understanding functional connectivity of human and model species. Multiple studies have shown that 2R-ohnologs (genes and proteins that have survived from the 2R-WGD) are enriched in signaling components relevant to developmental disorders and cancer. Our findings provide a link between AHR pathway’s evolutionary trajectory and its possible mechanistic involvement in pathogenesis.

## Introduction

The Aryl hydrocarbon receptor (AHR) is a transcription factor, originally identified for its role as sensor of dioxin (Okey, 2007), which belongs to the Per-ARNT-Sim (PAS) family of transcription factors, known for their role as sensor of environmental signals. PAS domains occur in a number of regulatory proteins, including “CLOCK” and ARNTL (Aryl hydrocarbon Receptor Nuclear Translocator-Like, also known as BMAL1: <u>B</u>rain and <u>M</u>uscle <u>A</u>RNT-<u>L</u>ike 1) proteins central to maintenance of circadian rhythms and “Hypoxia-inducible factors” (HIFs) involved in physiological adaptation to low oxygen (Gu et al., 2000; McIntosh et al., 2010).

Importance of this complex molecule has motivated research to understand its function and involvement in physiological response to environmental toxins (Avilla et al., 2020; Barroso et al., 2021; Bock, 2018; Gutiérrez-Vázquez and Quintana, 2018; Quintana, 2013; Quintana and Sherr, 2013). AHR is primarily activated by planar aromatic hydrocarbons (PAHs), and subsequently regulates the expression of some Cytochrome P450 (CYP) encoding genes and genes encoding conjugating and reducing enzymes (Jr., 1999; Nebert et al., 2004). The AHR genomic pathway is activated by ligand binding to the receptor followed by a conformational change which exposes the nuclear localization sequence (NLS) in the AHR N-terminal. This enables translocation of AHR to the nucleus where it undergoes dimerization with the aryl hydrocarbon nuclear translocator (ARNT). The AHR-ARNT heterodimer recognizes and binds to xenobiotic-response elements (XREs) in the genome and initiates transcription of target genes. AHR is primarily downregulated by AHR repressor (AHRR), which dimerizes with ARNT and competes for binding to AHR response elements (Evans et al., 2008; Hahn et al., 2009).

In the absence of a ligand, AHR is located in the cytoplasm for the majority of cell types in a complex with HSP90 (Heat Shock Protein 90), co-chaperone p23 and the aryl hydrocarbon receptor interacting protein, AIP. This enables AHR to maintain proper folding, recognition of ligand and subsequently, efficient translocation. In the absence of a ligand, AHR is unable to translocate to the nucleus and cannot interact with its main site of action; the DNA motifs of the promoter, the XREs of the target genes (Beischlag et al., 2008).

Recent studies have identified its role in development (Leung et al., 2017), which highlights its ancient origins and importance of understanding its evolution. Understanding evolutionary history of AHR and proteins involved in the AHR signal transduction pathway is necessary to advance our understanding on its function and assignment of cross-species functional connectivity between humans and model species.

AHR is an ancient protein (Mark E Hahn, 1998; Hahn et al., 2006, 2017). Although a significant amount of research has been carried out to understand the AHR signal transduction pathway and its role in health and disease (Barroso et al., 2021; Bock, 2018; Larigot et al., 2018; Rothhammer and Quintana, 2019), none of them to date has comprehensively studied its evolutionary origins. The vertebrate genome has been shaped by two rounds of whole-genome duplications (WGD) at the base of vertebrate evolution approximately 600 million years ago (Mya) (Dehal and Boore, 2005; Hokamp et al., 2003; McLysaght et al., 2002; Meyer and Schartl, 1999; Ohno, 1970), followed by lineage specific gene losses, which often complicate assignment of orthology.

Multiple possibilities regarding the number of ancient polyploidization events and their timings in relation to the cyclostome-gnathostome divergence have been proposed. (Summarized in Figure 1-(B), scenario1-3. Discussion on each model is beyond the scope of the present study. Number of detailed studies already cover this topic. Interested readers can refer to (Nakatani *et al*., 2021; Panopoulou and Poustka, 2005; Meyer and Schartl, 1999; Sidow, 1996; Dehal and Boore, 2005; Fried *et al*., 2003; Furlong *et al*., 2007; Mehta *et al*., 2013; Escriva *et al*., 2002; Kuraku *et al*., 2009; Smith *et al*., 2018)) This question is expected to evolve further with the increasing availability of high-quality genome assemblies in the future.

**Figure 1.**
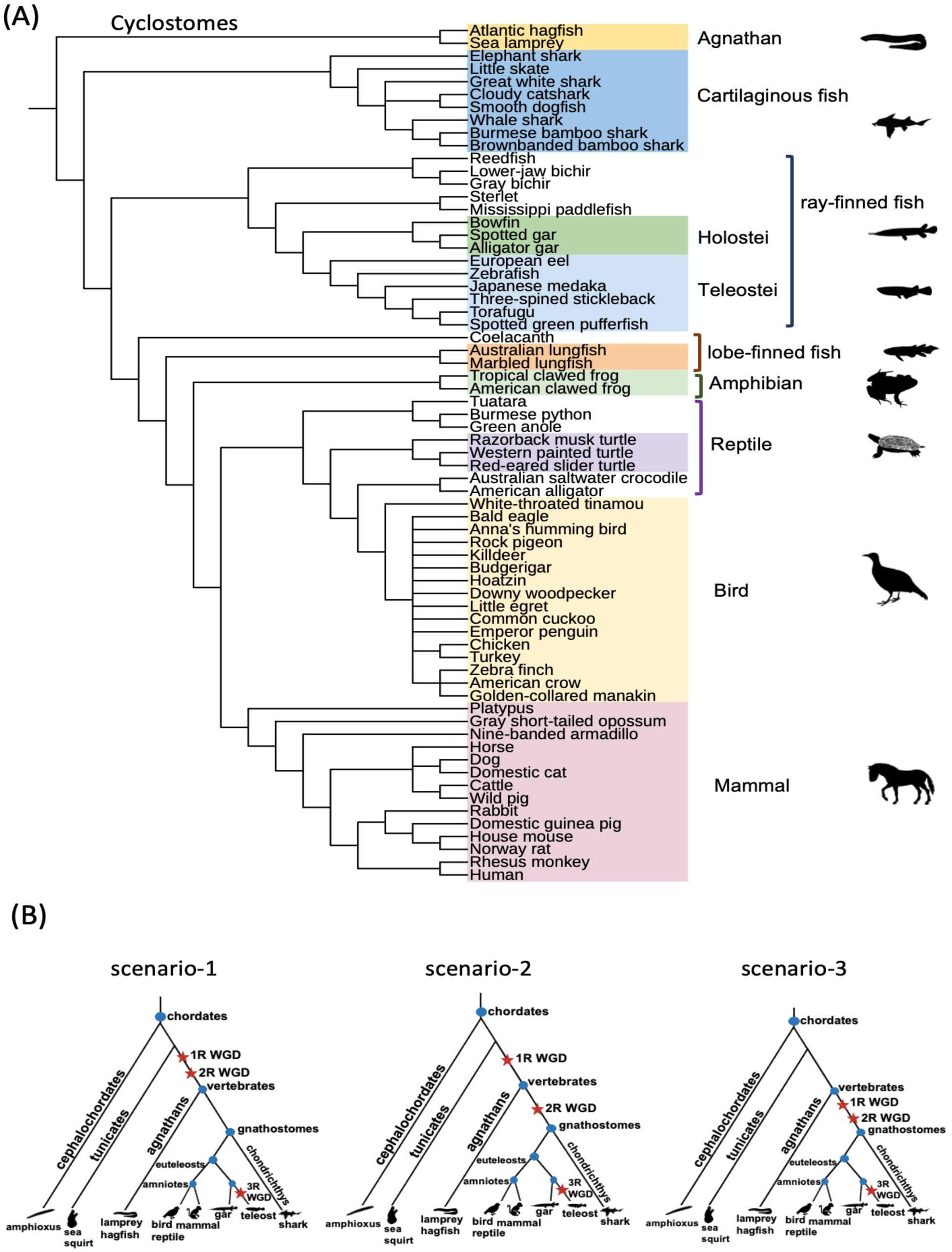
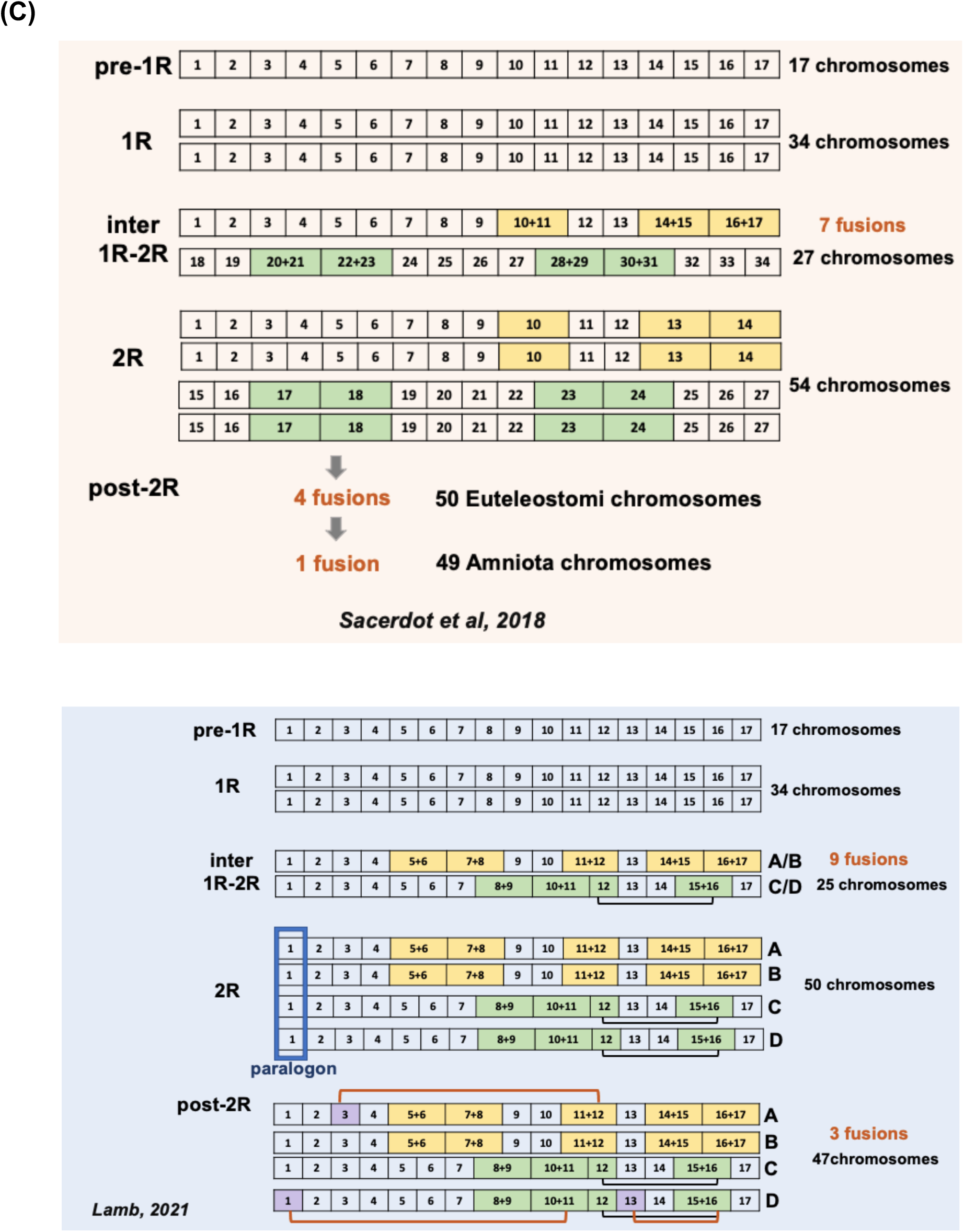
(A) Phylogeny of vertebrate species included in our study (B) Hypotheses regarding the timing of two rounds of WGD (C) Reconstruction of ancient chromosomes in two independent studies. (B) Multiple hypotheses have been proposed regarding the timing of two rounds of whole genome duplication (WGD) in relation to cyclostome-gnathostome divergence. Star denotes WGD. Scenario1: Some models predict two rounds of WGD occurred prior to cyclostome-gnathostome divergence. (Kuraku *et al*., 2009; Smith *et al*., 2013), scenario 2: Some models predict first round occurred before the split and the second round of WGD occurred after the split of cyclostomes and gnathostomes. (Escriva *et al*., 2002; Stadler *et al*., 2004), scenario 3: Some models predict that two rounds of WGD occurred after the cyclostome-gnathostome split in the proto-gnathostome lineage. (Mehta *et al*., 2013; Fried *et al*., 2003; Furlong *et al*., 2007). **(C) Two independent studies (Sacerdot *et al*., 2018; Lamb, 2021) recently reconstructed the pre-1R chromosome**. Both studies conclude that the pre-1R vertebrate karyotype consisted of 17 ancestral chromosomes. [above] pre-1R genome contained 17 chromosomes resulting in 34 chromosomes following 1R. Seven fusions occurred between 1R and 2R, leaving 27chromosomes. This resulted in 54 Vertebrata chromosomes by 2R, followed by four chromosome fusions leaving 50 chromosomes. An additional fusion led to the predicted Amniota chromosome comprising 49 chromosomes. Redrawn from *Secredot et al*., *2018*, Fig.5. Colored box indicate chromosome that underwent fusions. [below] According to *Lamb, 2021*, pre-1R genome contained 17 chromosomes leading to 34 chromosomes following 1R. Nine chromosomal fusions occurred after 1R WGD and prior to the second round of WGD. Colored boxes represent chromosomal fusions. According to the study, these 17 pre-1R chromosomes can be described as four “quadruplicates” A, B, C, D which each correspond to independent ohnologs that form the relationship ((A, B), (C, D)). Black line connecting chromosomes is the 9^th^ fusion (*not necessarily chronological). This left 25 chromosomes, resulting in 50 chromosomes in 2R (by doubling). Further fusions post-2R (orange lines, PQ=[1D, 10D+11D], PQ=[3A,11A+12A], PQ=[13D, 12D+15D+16D], P=paralogon, Q=quadruplicate) left a karyotype comprising 47 chromosomes prior to the radiation of bony-vertebrates. Blue rectangle represents paralogon (same for other chromosomes), each of these 17 paralogons correspond one-to-one with the 17 pre-1R ancestral chromosomes reported by *Sacerdot et al, 2018*. Redrawn from *Lamb, 2021* Fig.4.

While there are differing views on the timing and number of WGD, substantial amount of work supports the 2R-WGD hypothesis (Singh and Isambert, 2020; Sacerdot *et al*., 2018; Singh *et al*., 2015; Simakov *et al*., 2020; Lamb, 2021).

In addition to providing extra genetic material, 2R WGD has also contributed to the complexity of signaling pathways, particularly developmental pathways (Huminiecki and Heldin, 2010). Studies have shown that 2R-ohnologs (genes and proteins that have survived from the 2R-WGD) are enriched in signaling components relevant to developmental disorders and cancer (Singh *et al*., 2012; Tinti *et al*., 2014; Makino and McLysaght, 2010). One study showed that while 2R-ohnologs comprise only around 30% of protein-coding genes in the human genome, they carry 42-60% of the somatic mutations in transcript-coding genes in 30 types of cancers that were examined (Tinti *et al*., 2014).

Since functionally related genes and proteins evolve together (Goh *et al*., 2000), understanding the evolutionary trajectory of the AHR signaling pathway is important for the proper interpretation of comparative analysis, especially when using model organisms to study human gene functions. The evolutionary origin of the AHR gene family probably predates the origin of stem bilaterians (Hahn et al., 2017), but the ability of the Ahr homologs of basally diverging bilaterians to respond to environmental toxins remains unknown.

Understanding the evolutionary history of AHR and proteins involved in the AHR signal transduction pathway can provide a broader perspective on their function that will complement the results of experimental studies. In order to distinguish orthologs from ancient non-orthologous homologs, it is crucial to understand evolutionary origins of this transcription factor and its partners. Additionally, as noted by earlier studies on AHR (Mark E. Hahn, 1998), this can assist in filling knowledge gaps in understudied taxonomic groups. In this study, we examined the evolutionary history of the AHR gene family and proteins involved in the AHR signal transduction pathway, AHR, AHRR, ARNT, ARNT2 and AIP. Understanding the evolutionary history of this gene family forms the basis of assigning functional orthology and is crucial for translational research. Here, we show how multiple rounds of whole genome duplications shaped the AHR pathway genes.

## Methods

### 1. Phylogenetic analysis of AHR proteins

Human protein sequences belonging to the AHR pathway were downloaded from UniProtKB database. Homologous sequences were extracted in a first round by local blastp searches in the STRING database (protein.sequences.v11.5.fa) (Szklarczyk et al., 2019) using each AHR pathway protein as the query. Multiple sequence alignments (MSA) were constructed for each query protein and homologs by MAFFT (with option FFT-NS-2) (Katoh et al., 2002; Katoh and Standley, 2013). HMM models were made from MSAs for each of these, followed by profile HMM search (option: hmmsearch after hmmbuild) with HMMER v3.2.1 (www.hmmer.org) in the STRING protein database for a second round of homology search. Profile HMM based methods are amongst the most successful procedures for detecting remote homology between proteins (Madera and Gough, 2002). Gblocks 0.91b (Castresana, 2000) was used to remove ambiguously aligned blocks. This Multiple Sequence Alignment was then filtered to a list of 90 eukaryotic species (Supplementary material) to only contain sequences of these species. The selection of species is based on the estimated evolutionary origin of the Per-ARNT-Sim pathway described in (Leung et al., 2017) and an even spread of species over the taxonomy. This was done prior to phylogeny construction to shorten the computational time and to minimize short, ambiguous branches. Maximum likelihood trees were constructed by RAxML-NG v.1.1(Kozlov et al., 2019) using the LG+FC+G8m model with 200 bootstrap replicates. This gene tree was then reconciled with a species tree (NCBI species tree) using NOTUNG version 2.9.1.5 (Chen et al., 2000) to infer gene duplication and loss events (Szöllősi et al., 2015) followed by rearrangement with default edge weight threshold of 90%. Rearrangement was used to determine the tree with a minimal event score (Stolzer et al., 2012). Preprocessing of trees was done by ETE3 (Huerta-Cepas et al., 2016) and taxonkit (Shen and Ren, 2021), and were drawn in iTOL (Letunic and Bork, 2021).

### 2. Assessment of conserved synteny

Synteny analysis was carried out using Synteny Database (Catchen et al., 2009). The Synteny Database is designed to compare genomes that have undergone one or more whole-genome duplications and can detect chromosome inversions and translocations, as well as ohnologs gone-missing of individual gene families. This approach complements the use of BLAST and phylogenetic reconstructions and provides additional evidence to infer the evolutionary history of the gene family independent of sequence identities. Additionally, Genomicus (Muffato *et al*., 2010) v93.01 was used to study synteny in detail.

## Results

### 1.1 Phylogenetic analysis of the AHR gene family

Human proteins of the AHR pathway were subjected to two rounds of homology searches to extract remote homologs, followed by construction of Multiple Sequence Alignments and phylogenetic trees for each member of the pathway. To understand the history of gene gain and loss in the AHR family, it is important to first understand the phylogeny of the family members. We inferred the gene histories by reconciliation of the species tree with the gene trees.

The reconciled phylogeny of the AHR protein shows that AHR and AHRR emerged by an ancient duplication (duplication A, Figure 2) in gnathostomes followed by a major duplication (duplication G) leading to clades AHRR and AHR2. Full-length homologs of *Drosophila* single-minded protein *ss*, urochordate *Ciona*, and agnathan lamprey PAS domain-containing protein are present prior to this duplication which suggests the ancestral AHR gene was already present in the common ancestor of these species. The *C. elegans* genome also contains an *ahr* gene, but the *ahr-1* gene is not present in the tree due to the sequence similarity being too low. The AHR clade (highlighted in yellow) contains homologs of cartilaginous, fish cloudy catshark, bony fishes, reed fish and *Latimeria* in addition to teleost fishes. This clade contains four duplications; a teleost specific duplication B, a lineage specific duplication in American clawed frog (duplication C), duplication D in some reptiles (western painted turtle, American alligator and Australian saltwater crocodile), and a duplication (duplication E) in the bird lineage, in chicken, bald eagle, rock pigeon and Passeriformes zebra finch and golden-collared manakin.

**Figure 2.**
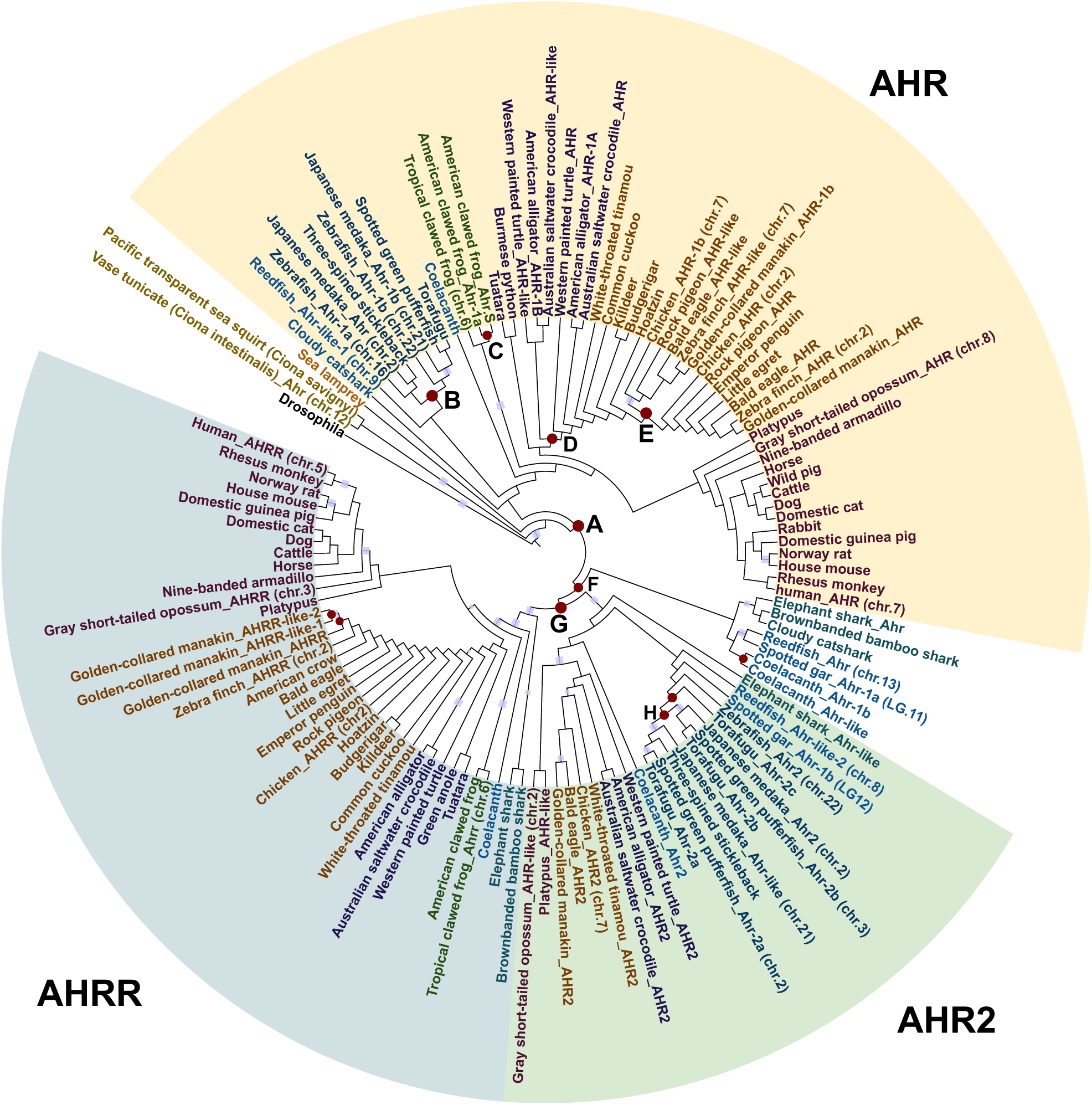
Phylogenetic tree of AHR gene family. Phylogenetic tree of the AHR gene family shows ancient duplication in the gnathostome lineage (duplication A) led to two distinct clades of AHR and a clade consisting mostly of AHRR and AHR2. Brown dots are duplication nodes. Blue rectangles are edges with bootstrap value above 90.

Among fish species, elephant shark, along with coelacanth and spotted gar, are known to have slow-evolving genomes (Venkatesh *et al*., 2014; Braasch *et al*., 2015, 2016; Amemiya *et al*., 2010). The elephant shark Ahr, spotted gar Ahr (LG11) and Reed fish Ahr are located in an independent clade resulting from duplication F, along with other homologs. This may have resulted as a separate clade due to these slow evolving genomes preserving their sequence and not evolving as fast as the sequences in the AHR clade.

While most mammals including humans have two homologs, AHR and AHRR, some reptiles, birds, and fish species have three or more Ahr homologs (Figure 2).

The AHRR clade contains homologs of cartilaginous fish, *Latimeria*, amphibians, some reptiles, birds and mammals. Two lineage specific duplications are observed in the bird, golden-collared manakin.

The AHR2 clade contains two duplications in the teleost lineage. This clade contains cartilaginous fish, bony fish, reptiles, birds and mammals; platypus and opossum.

To test historical relationships among AHR paralogs, we examined a data set independent of AHR gene sequence by comparative genomic analysis of the genomic neighborhoods surrounding AHR genes in the genomes of fish and other vertebrates. The duplication B in the AHR clade and duplication H in the AHR2 clade consist only from teleost species. Thus, these duplications may be the third round of whole genome duplication (3R WGD) which occurred at the base of teleost radiation.

### 1.2 Analysis of conserved syntenies for the AHR gene family

As an independent line of evidence for each duplication event, we carried out a synteny analysis to distinguish orthologs from distant non-orthologous homologs.

The slow-evolving spotted gar genome was used as a reference to infer teleost-specific duplications and as a bridge between human and teleost species. The spotted gar genome shows low rates of protein evolution and extremely low rates of chromosome evolution (Braasch et al., 2016). Despite teleost species such as the zebrafish, fugu, and medaka commonly used as model species in research, assignment of true orthology is complicated due to the third round of whole-genome duplication (3R WGD) that occurred at the base of teleost radiation (Figure 1-(B)) followed by lineage-specific gene losses in these lineages (Meyer and Schartl, 1999; Nakatani et al., 2007; Postlethwait et al., 2004; Taylor et al., 2003; Wittbrodt et al., 1998). Recent studies have suggested that using spotted gar as reference can reveal features between teleost and humans that are not evident in direct comparisons (Parichy, 2016) and can be particularly useful in understanding the evolutionary and pathological significance of gene regulatory variation (Braasch et al., 2015).

### 1.3 Analysis of genes in medaka supports the hypothesis that ahr1b and ahr2 are co-orthologs to spotted gar ahr2

Given that there are two sequences for most of the species resulting duplication, it is possible that these duplications (duplication B and H) represent teleost specific 3R WGD. If that is the case, that is, if teleost paralogs under clades of duplication B and duplication H resulted from 3R WGD, two homologs of each teleost species (in-paralogs) would be co-orthologous to spotted gar *ahr* (that did not undergo 3R). The homologs under clades from these duplications, medaka *ahr-1b* (8090.ENSORLP00000022794) and *ahr-like* (8090.ENSORLP00000022790) are located on chromosome 21 (Ola21), while *ahr* (8090.ENSORLP00000000168) and *ahr2* (8090.ENSORLP00000000166) are located on chromosome 2 (Ola2). To see if these two duplications (duplication B in the AHR clade and duplication H in the AHR2 clade) represent teleost specific 3R WGD, we checked for conserved synteny between medaka and spotted gar. Orthology dot plot shows a high degree of conservation between spotted gar chromosome LG12 and medaka Ola 2, Ola21(Figure 3-(A)) spanning through the entire chromosome, indicative of resulting from large-scale duplications, which suggests multiple regions on these chromosomes are likely to be paralogons. (A paralogon is a set of paralogous chromosomal regions, derived from a common ancestral region.)

**Figure 3.**
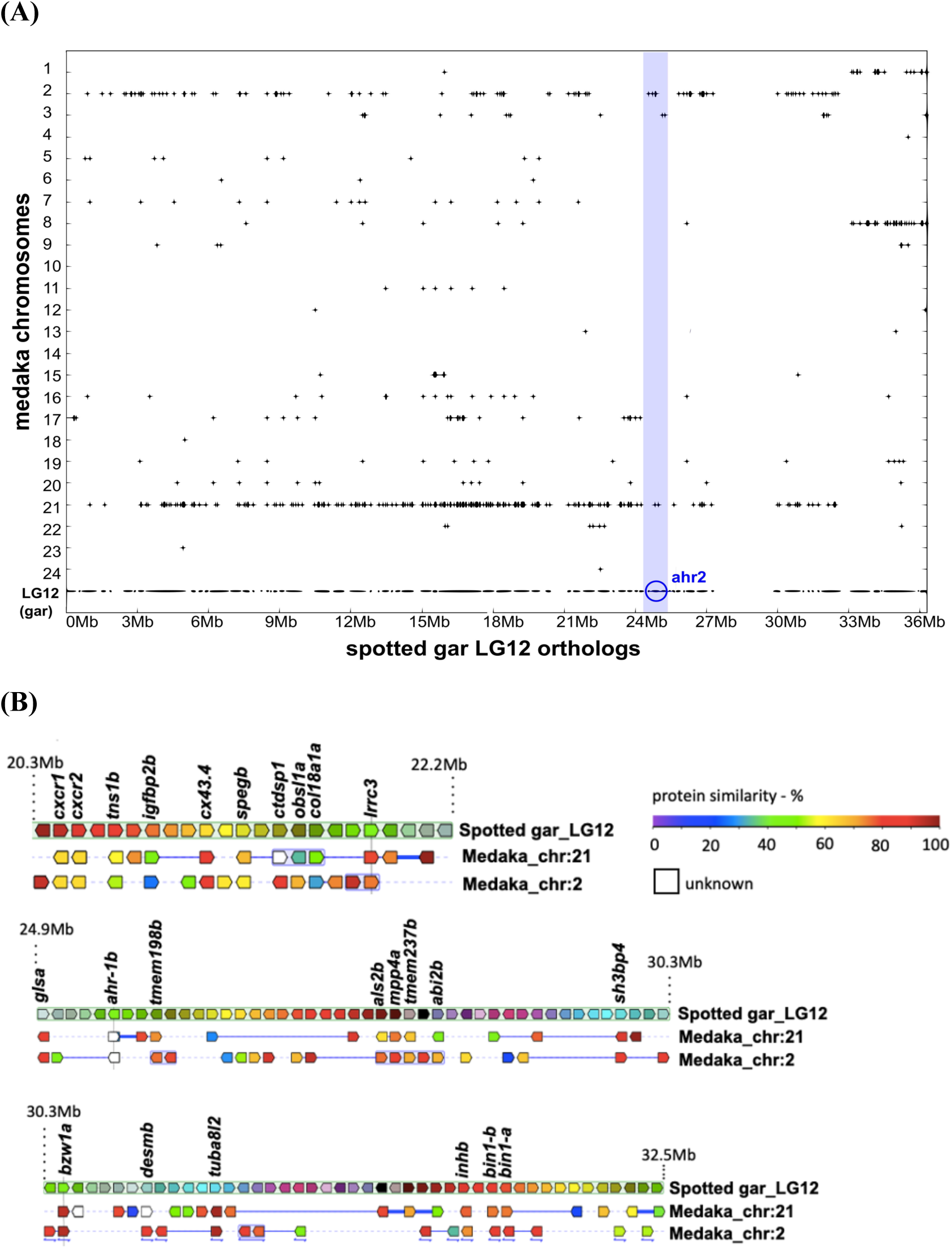
Conserved synteny on the ahr-1b chromosomal region. **(A)** Orthology dot plot of orthologous genes between spotted gar chromosome LG12 and medaka chromosomes. X-axis shows genes on spotted gar chromosome LG12 and y-axis are the medaka chromosomes. Plus (+) indicates that the orthologous gene is present on the medaka chromosome. Order of genes on medaka chromosome don’t necessarily represent actual location on its chromosome. Location of ahr-1b gene on spotted gar chromosome LG12 is shown in blue circle. Spotted gar LG12 and Ola 2, Ola21 show high degree of conservation. Dot plot generated by Synteny Database. **(B)** Co-duplicated genes (labeled genes) present on chromosomal region surrounding ahr-1b. Location (Mb) indicates chromosomal region on spotted gar genome. The genes on either side of the reference gene are shown in correct order and orientation. Orthologs of the reference gene and its flanking genes are shown below the reference chromosome, organized by species and chromosome. Colors of genes represent percentage of sequence identity between homologs (with gene on the reference chromosome). Consistent with the database, double-headed arrow under a block of genes indicates that the order of genes shown was flipped around. Thick blue line between genes is equivalent to a “gap” in the alignment of this species and thin blue line corresponds to a “break” of the alignment (two genes are linked in order, but at least one gene separates them in other species). Dotted line means no alignment. Synteny analysis was performed with Genomicus (Muffato et al., 2010) version 93.01 with the Alignview tool.

To confirm if the duplication between medaka homologs on Ola2 (*ahr2, ahr*) and Ola21 (*ahr-1b, ahr-like*) occurred from 3R WGD, the following would confirm (or reject) this hypothesis. If the two medaka paralogs on chromosome Ola21 (*ahr-like*: 8090.ENSORLP00000022790 and *ahr-1b*: 8090.ENSORLP00000022794) emerged as a result of 3R WGD, first, these two homologs would show a greater degree of syntenic conservation compared to solely by chance (sufficient number of genes surrounding these two sequences will be paralogous to their counterpart occurring from the ancestral pre-3R chromosome, Ola2). Second, these neighboring paralogs will be orthologous to the neighbors of spotted gar *ahr-1b*.

Our results (Figure 3-(A)) show strong conservation spanning over 25Mb (25 megabases) in the area around *ahr* homologs on Ola2 and Ola21. This highly conserved genomic neighborhood provides evidence that this region of Ola2/Ola21 was produced in a large-scale duplication event. Chromosomal neighborhood surrounding spotted gar *ahr-1b* is shown in Figure 3-(B). Multiple co-duplicating genes can be observed in this region.

### 1.4 Vertebrate Ahr homologs emerged from an ancient chromosome prior to 1R — a pre-1R paralogon

The above findings raise questions if traces of ancient chromosome duplications are present in other species. To address this, we checked the chromosomal location of homologs in each evolutionarily representative lineage of vertebrate species present in our phylogeny. The results are summarized in Supplementary Table 1. This revealed that these genes fall on ancient pre-1R chromosomes, which, after WGD, resulted in ohnologs (gene duplicates originated by genome duplication).

Two recent studies reconstructed the ancient vertebrate karyotype preceding the first round of WGD (1R) in the stem vertebrates by different approaches (Figure 1-(C)). Both of these studies concluded that vertebrate karyotype arose from an ancestral chordate karyotype which comprised 17 chromosomes (Lamb, 2021; Sacerdot et al., 2018).

According to (Sacerdot et al., 2018), ancestral gene of AHR and AHRR was located on pre-1R chromosome 1(for details, see (Sacerdot et al., 2018)).

For each species AHR homolog from our phylogeny identified above, its chromosomal locations fall on paralogon-1 (ancient chromosome 1), ohnolog-D according to (Lamb, 2021) (for details, see (Lamb, 2021) FIG 1. PQ = 1D). These findings, combined with the fact that *Ahr* homolog is present in the urochordate *Ciona* (Figure 2), provides additional support that *ahr* ancestral gene was already present prior to two rounds of WGD at the stem of vertebrate evolution.

### 2.1 Phylogenetic analysis of ARNT gene family

Phylogenetic analysis of the ARNT gene family reveals an ancient duplication in gnathostomes (duplication A) leading to a clades ARNT and ARNT2, and another duplication (duplication B) that diverged into ARNTL1 and ARNTL2 clades (Figure 4). Full-length homologs of Urochordate *Ciona, C*.*elegans aha-1*, and *Drosophila tgo* are present in the phylogeny as well, consistent with previous studies (Emmons et al., 1999; Qin and Powell-Coffman, 2004; Sonnenfeld et al., 1997). Cartilaginous fish homologs are present in clades except ARNTL2, in addition to spotted gar, bony-fishes, teleost, amphibians, reptiles, birds and mammal species homologs, which confirms this gene family was preserved in these evolutionary lineages. The close relationship of ARNT genes with the circadian genes ARNTL1 and ARNTL2 highlights the importance of these genes and ancient origins of the AHR signal transduction pathway.

**Figure 4.**
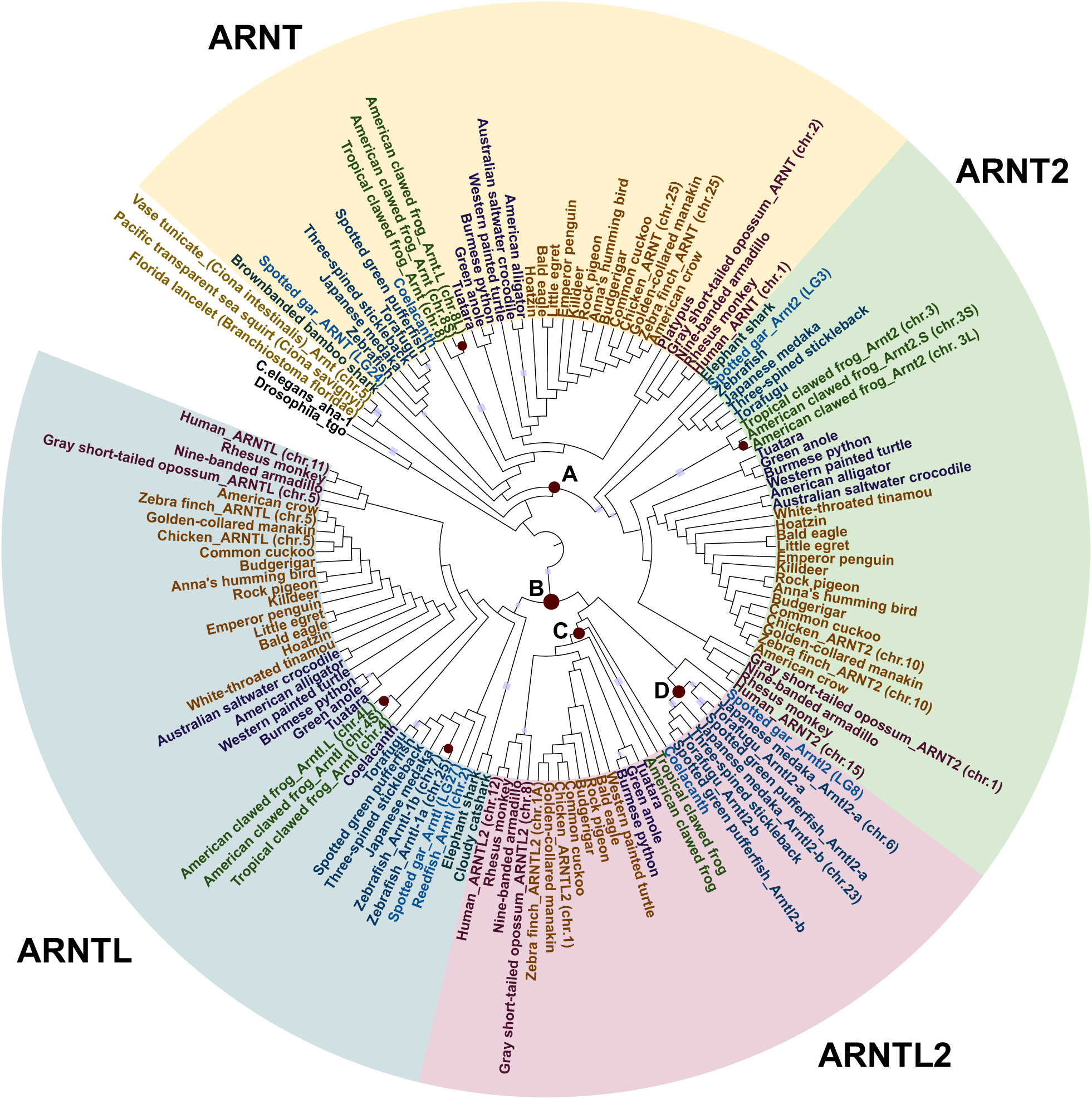
Phylogenetic tree of ARNT gene family. Phylogenetic tree of the ARNT gene family proteins shows ancient duplication in gnathostomes leading to two major duplications of ancient ARNT/ARNT2 and ARNTL/ARNTL2. Brown dots are duplication nodes. Blue rectangles are edges with bootstrap value above 90.

The ARNT, ARNT2 and ARNTL clades contain three lineage specific duplications in American clawed frog. Additionally, the ARNTL2 clade contains a duplication (duplication D) which consists only from teleost.

### 2.2 Analysis of conserved syntenies in the ARNT gene family

Synteny analysis of the ARNT gene family shows a high degree of conservation between human ARNT genes and spotted gar homologs (Supplementary Figures 1-4). This indicates that a large part of the human chromosomal region surrounding ARNT genes has likely resulted from ancient paralogons. These findings are consistent with recent studies which have mapped this gene family’s location on pre-1R chromosomes (Sacerdot et al., 2018), (Supplementary Table1).

Supplementary figure 1 shows orthologs between spotted gar chromosome LG24 and human chromosomes. Hsa1, where ARNT is located, shows syntenic conservation with LG24 (location of spotted gar ARNT gene).

Supplementary figure 2 shows spotted gar LG3 orthologs with human chromosomes. LG3 shows strong syntenic conservation with HSa1, HSa11, HSa15 and weak conservation with Hsa3, Hsa6 and HSa13. Human chromosome 15, where ARNT2 gene is located, is known as one of the chromosomes (along with HSa14) almost entirely composed of genes from a single pre-1R chromosome (Sacerdot et al., 2018). Strong syntenic conservation spanning over the entire length of chromosome can be observed between LG27 and HSa 11, surrounding the ARNTL neighborhood (Supplementary figure 3). This supports the findings that this gene in fact, resulted from an ancient paralogon. Supplementary figure 4 shows orthologous genes between spotted gar chromosome LG8 and human chromosomes. Strong syntenic conservation can be confirmed between LG8 and HSa12, HSa7. These two chromosomes, HSa12, HSa7 along with HSa2 and HSa17 are Hox-bearing chromosomes, frequently used as evidence for 2R (Panopoulou and Poustka, 2005). These findings confirm the ancient origins of the ARNT gene family.

### 2.3 Analysis of arntl2 genes in medaka supports the hypothesis that these are co-orthologs to spotted gar arntl2

The ARNTL2 clade in the phylogeny contains a duplication that only contains teleost species homologs, which raises the question if this duplication represents teleost WGD, 3R.

To confirm this, the following would confirm (or reject) this hypothesis. 1) These medaka *arntl2* homologs (*Arntl2-a* _chr.6 and *Arntl2-b*_chr.23) will be paralogous (and show a greater degree of syntenic conservation than expected by chance) and 2) these homologs would share a genomic neighborhood surrounding arntl2 higher than expected by chance with species that did not undergo 3R. Medaka *arntl2* is located on Ola6 (ENSORLP00000013730) and Ola23 (ENSORLP00000011958). Synteny analysis in the genomic neighborhood of *arntl2* homolog shows strong conservation between spotted gar LG8 and medaka Ola6, 23 (Figure 5A), which confirms this hypothesis. The high degree of conservation (synteny) between teleost homologs and medaka *arntl2* (Figure 5B,5C,5D) further supports the hypothesis that this duplication represents 3R.

**Figure 5.**
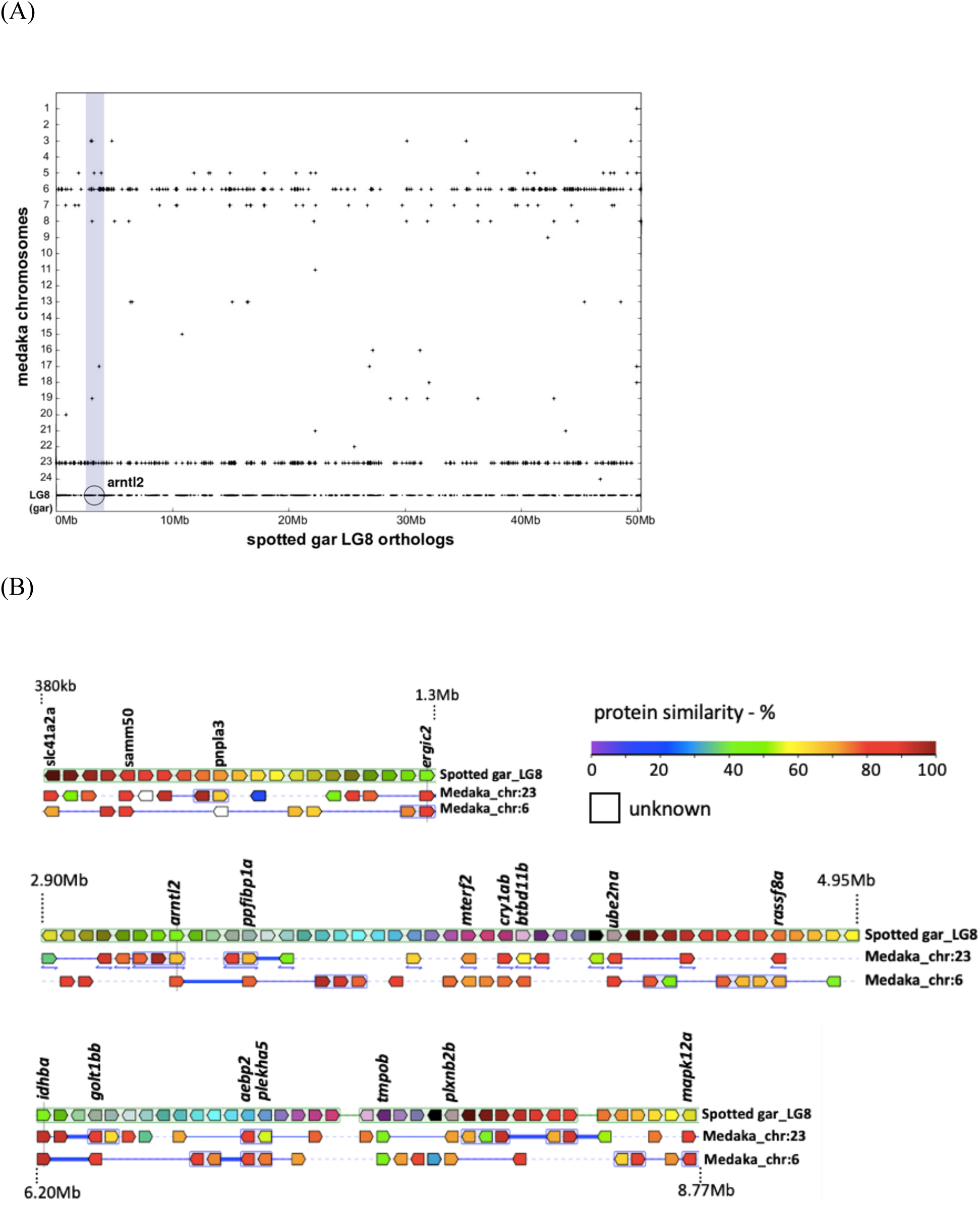
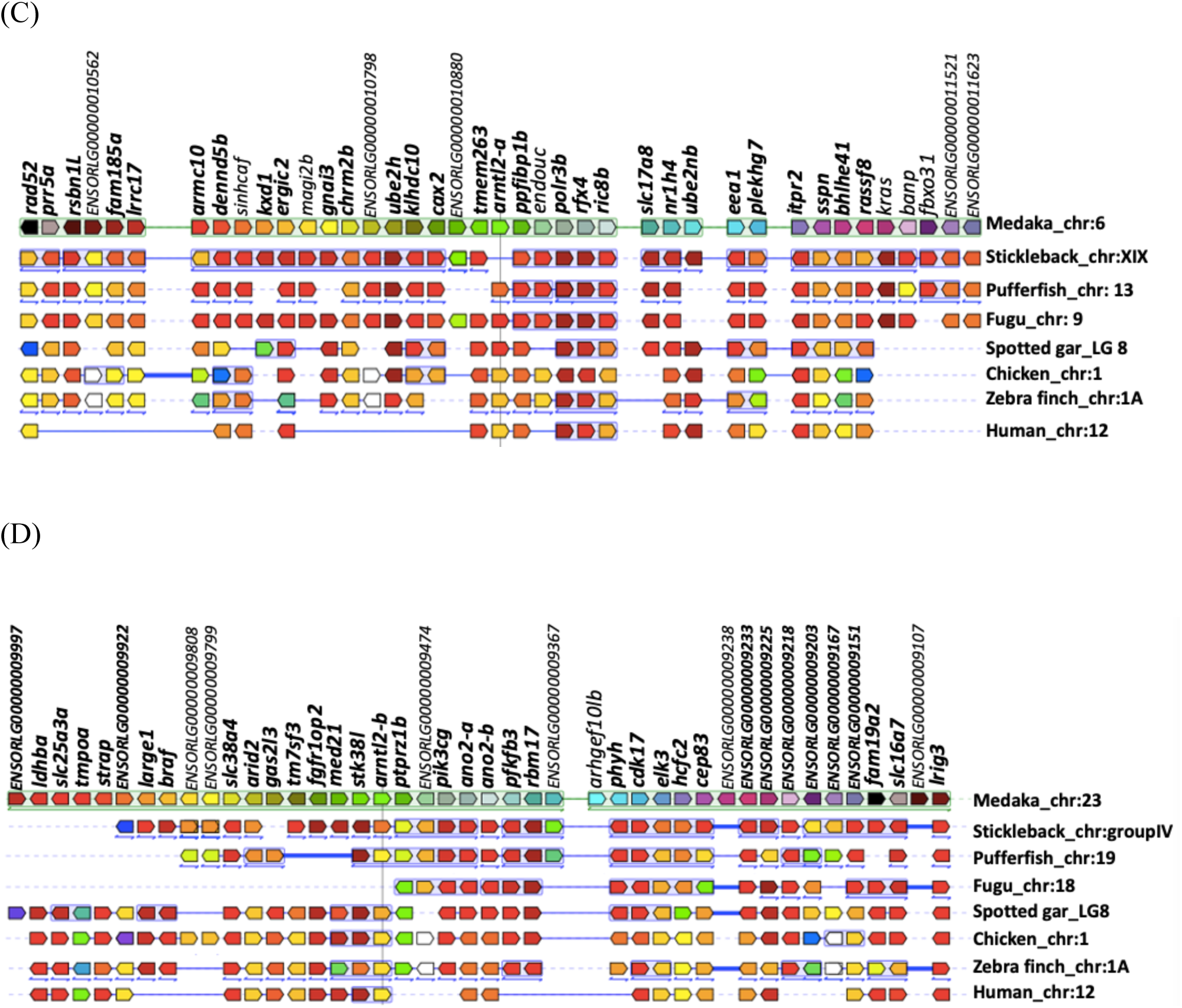
Conserved synteny on the arntl2 chromosomal region. **(A)** Dot plot of spotted gar LG8 genes and medaka chromosomes. Strong conservation between LG8 and Ola6, Ola23 can be confirmed. **(B)** Co-duplicated genes present on chromosomal region surrounding the arntl2 gene (vertical line). Chromosomal location (Mb) on the spotted gar genome is shown. Synteny analysis was performed with Genomicus (Muffato et al., 2010) version 93.01 with the Alignview tool. **(C)** Conserved synteny surrounding medaka arntl2-a gene on chromosome Ola6. Gene names in bold indicate genes present on both medaka Ola6 and spotted gar LG8. **(D)** Conserved synteny surrounding medaka arntl2-b gene (vertical line) on chromosome Ola23. Gene names in bold indicate genes present on both medaka Ola23 and spotted gar LG8. All these chromosomes fall on the same pre-1R paralogon described by (Lamb, 2021). The genes on either side of the reference gene are shown in correct order and orientation. Orthologs of the reference gene and its flanking genes are shown below the reference chromosome, organized by species and chromosome. Colors of genes represent percentage of sequence identity between homologs (with gene on the reference chromosome). Consistent with the database, double-headed arrow under a block of genes indicates that the order of genes shown was flipped around. Thick blue line between genes is equivalent to a “gap” in the alignment of this species and thin blue line corresponds to a “break” of the alignment (two genes are linked in order, but at least one gene separates them in other species). Dotted line means no alignment.

### 2.4 Vertebrate Arnt homologs emerged from an ancient chromosome prior to 1R — a pre-1R paralogon

As described for AHR homologs, chromosomal location of vertebrate ARNT gene family homologs present in four major clades in the phylogeny are listed in Supplementary Table 1. These findings strongly support ARNT and ARNTL ancestral genes emerged from pre-1R ancestral genes prior to the first round of whole-genome duplication that occurred at the stem of vertebrate evolution. In fact, in a recent study by (Sacerdot et al., 2018), where they reconstructed ancient pre-1R chromosomes, show ARNT/ARNT2 ancestral gene on pre-1R chromosome 7 and ARNTL/ARNTL2 ancestral gene on pre-1R chromosome 6 (for details, see (Sacerdot et al., 2018) supplementary file 8). Additionally, the phylogeny contains homologs of lancelet (*Branchiostoma floridae)* and *Ciona* which further support these findings.

### 3.1 Phylogenetic analysis of the AIP gene family

Phylogenetic analysis of AIP protein revealed AIP and AIPL (Aryl hydrocarbon receptor Interacting Protein Like) emerged from an ancient duplication in gnathostomes (Figure 6, duplication A). Full-length homologs of *C*.*elegans aipr-1* and *Drosophila CG1847* are also present in the phylogeny suggesting ancestral *aip* gene already existed in the bilaterian ancestor.

**Figure 6.**
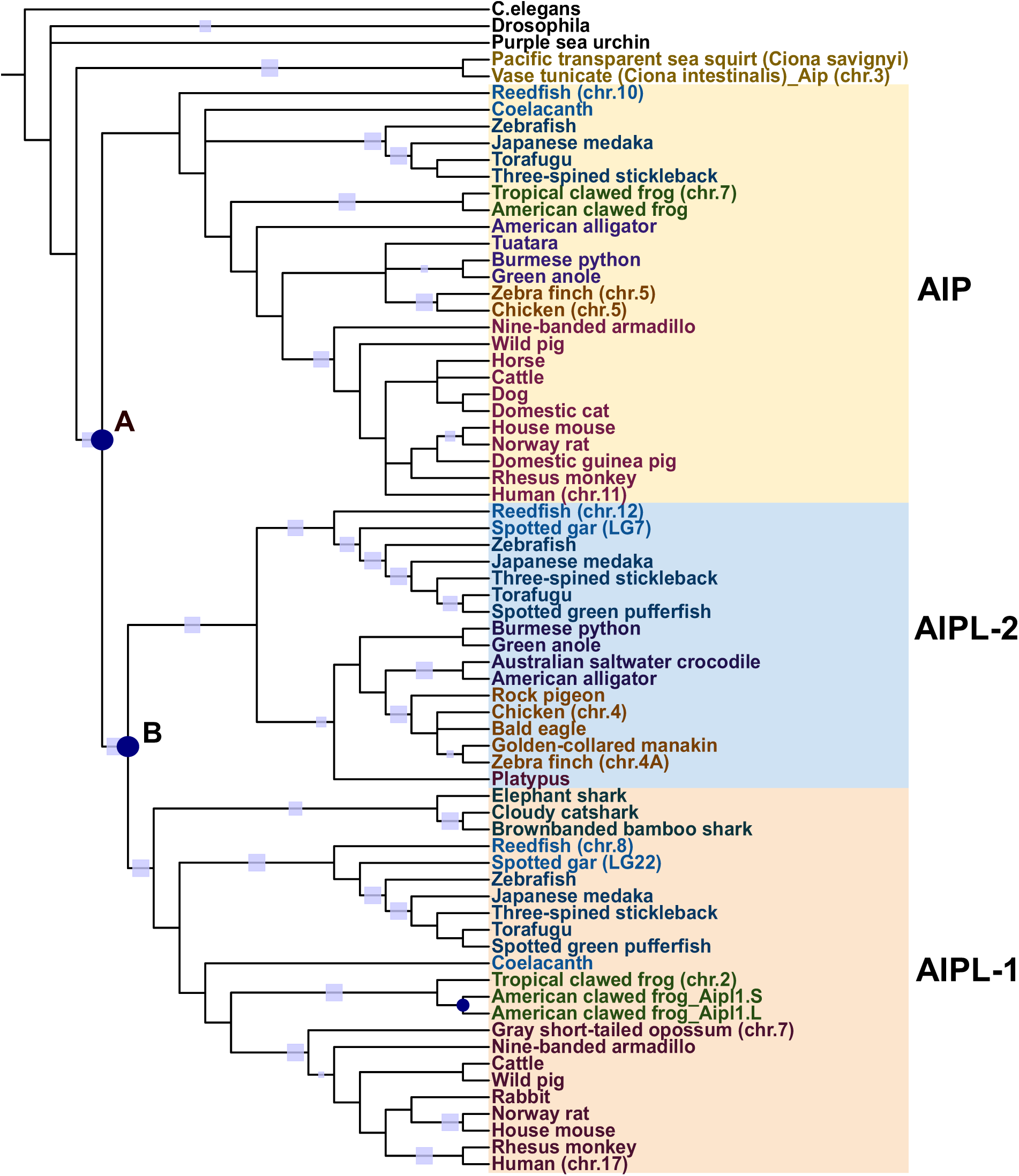
Phylogenetic tree of AIP gene family. Phylogenetic tree of the AIP gene family proteins shows ancient duplication in gnathostomes leading to two major clades of AIP and AIPL. Blue dots are duplication nodes. Light blue rectangles are edges with bootstrap value above 90.

The AIP clade contains homologs of bony fish, amphibians, reptiles, birds and mammals. The phylogeny also revealed that vertebrate-specific AIPL proteins fall into two main clades: (1) clade consisting mainly of AIPL-1 and (2) AIPL-2 resulting from duplication B. The AIPL-1 clade contains a lineage specific duplication in American clawed frog. While the AIPL-1 clade contains a distinct clade of cartilaginous fish, homologs of bony fish, amphibians and mammals; homologs of bird species, reptiles and the egg-laying mammal platypus are only found in the AIPL-2 clade. Given that cartilaginous fish homologs and slow-evolving genomes, elephant shark and spotted gar, are absent from AIP clade, it is likely that AIPL is the ancient form of this gene family. The fact that reed fish contains two homologs of aipl suggests additional copies of aipl in teleost haven’t necessarily resulted from teleost specific duplication events, but rather lineages with less than two homologs of aipl may have undergone gene loss of this homolog.

### 3.2 Analysis of conserved syntenies for the AIP gene family

Synteny analysis of spotted gar *aipl-1* and medaka *aipl-1* revealed a highly conserved region spanning over 2Mb surrounding *aipl-1* homologs on LG22 and Ola13 (Figure 7A). This spotted gar chromosome LG22 is thought to be part of pre-1R ancestral chromosome (details described in the next section) (Lamb, 2021), which supports the ancient origins of this gene. Additionally, the presence of *aip* homolog in the urochordate *Ciona* suggests this gene was already present at the base of vertebrate evolution.

**Figure 7.**
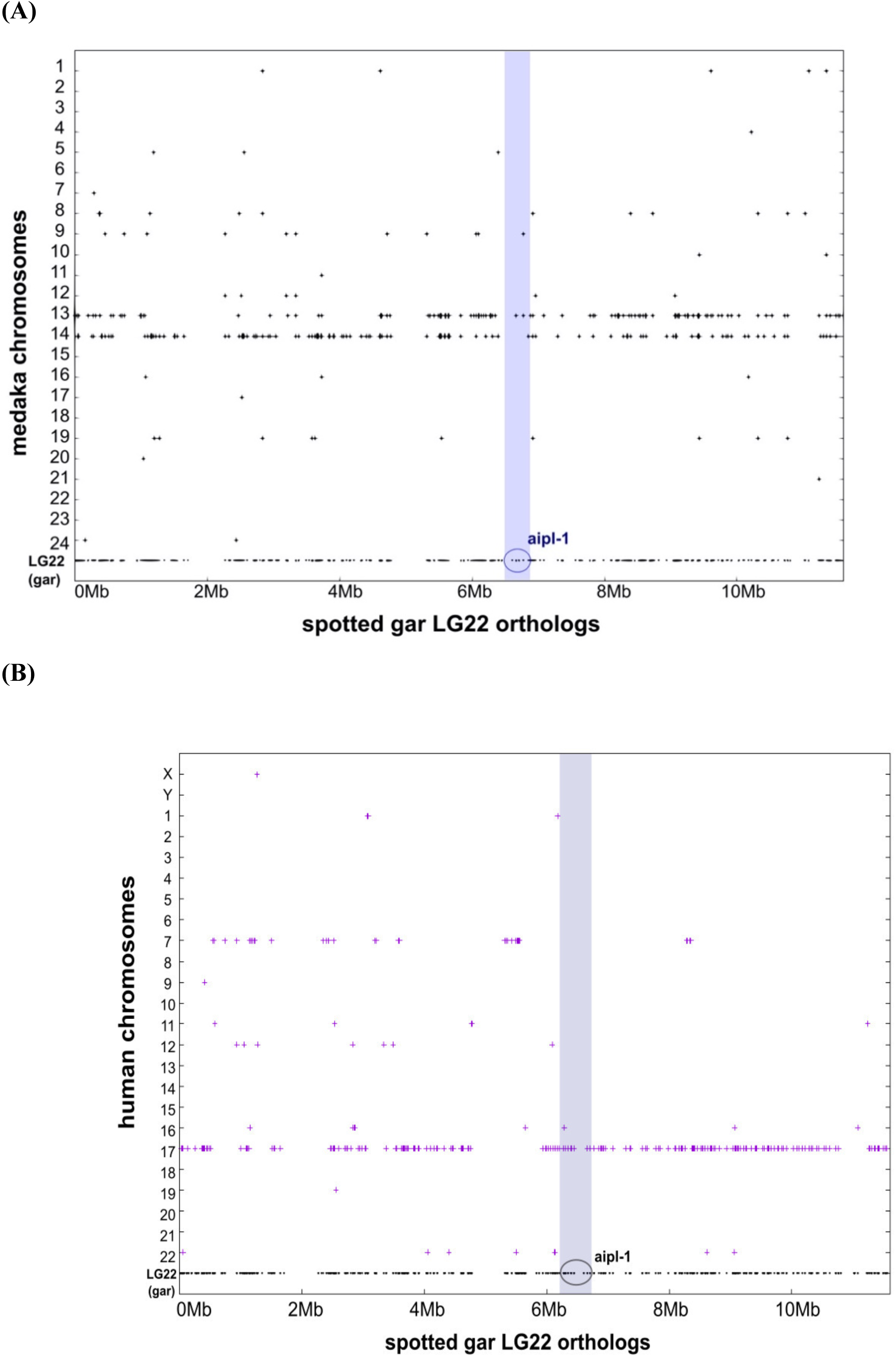
Conserved orthology in AIPL-1 evolution. **(A) Orthology dot plot showing the medaka orthologs of spotted gar LG22 genes**. Medaka orthologs of LG22 are plotted vertically above the corresponding dot on their respective medaka chromosome. LG22 shows strong orthology with Ola13 and Ola14. **(B) Orthology dot plot showing the human orthologs of spotted gar LG22 genes**. Human orthologs of LG22 are plotted vertically above the corresponding dot on their respective human chromosome. LG22 shows strong orthology with Hsa17 and weaker orthology with Hsa7. High degree of conservation along the entire length of LG22 and Hsa17 confirms that these occurred from a large-scale duplication event. Location of genes on LG 22 represent actual location of these on the chromosome, while human genes (orthologs to spotted gar LG22) on Hsa chromosomes do not necessarily represent their actual chromosomal location. Orthology dot plot created by Synteny Database.

### 3.3 Vertebrate AIP homologs are conserved on ancient pre-1R paralogon

Chromosomal locations of vertebrate *aip* homologs in the phylogeny (summarized in Supplementary Table 1) point to a possibility that these (*aip, aipl-1, aipl-2)* originated from ancestral paralogon as described in recent studies (Figure 1-(C)). Our findings on chromosomal locations of the homologs supports that *aip* homologs are 2R-ohnologs. High level of conservation along the spotted gar LG22 and human chromosome 17 revealed by orthology dot plot confirms that *aipl-1* emerged from a large-scale duplication event (Figure 7B).

Chromosomal location of opossum *AIPL-1* (chr.7) under AIPL-1 clade and zebra finch, chicken (chr.19, for both) show synteny with human chromosome 3 and chromosome 17 (Supplementary Figure 5). This is not surprising as it is thought that chromosomal rearrangements took place in the eutherian ancestor as described in (Robinson and Ruiz-Herrera, 2008).

### 3.4 Aipl-2 Ohnolog gone-missing in mammals and amphibians

Multiple studies have shown very slow rates of genome evolution in Holostei (gar and bowfin) and sauropsids (birds and reptiles) (Braasch et al., 2016; Shaffer et al., 2013). Our findings that *aipl-1* ancestral gene falls on an ancient pre-1R paralogon raise questions if the other *aipl* homologs under AIPL-2 clade in our phylogeny (Figure 6) have also resulted from an ancient WGD event. Chromosomal location of the spotted gar, chicken, zebra finch, and reed fish homologs under this AIPL-2 clade all fall on the same paralogon, C12,15 or 16 (these are fused, thus identical) based on (Lamb, 2021) (Supplementary Table1, Figure 10, Figure 1-(C)).

Synteny analysis with chicken AIPL homolog as reference shows strong conservation with human, rhesus, rat, mouse, and opossum X chromosomes. Additionally, frog chromosome 8 also shows syntenic conservation with this region (Figure 8). These findings all point that homologs in AIPL-2 clade in our phylogeny (Figure 6) share an ancestral *aipl-2* gene on the pre-1R chromosome. Strong syntenic conservation along the chromosomal region surrounding spotted gar *aipl-2* (LG7) and human chromosome X support this finding (Figure 9). Therefore, this region on human, rhesus, mouse, rat, opossum and frog chromosomes (between genes *BTK* and *DRP2*) is likely the location of the ancient ohnolog gone-missing (ogm) *aipl-2* gene in these lineages.

**Figure 8.**
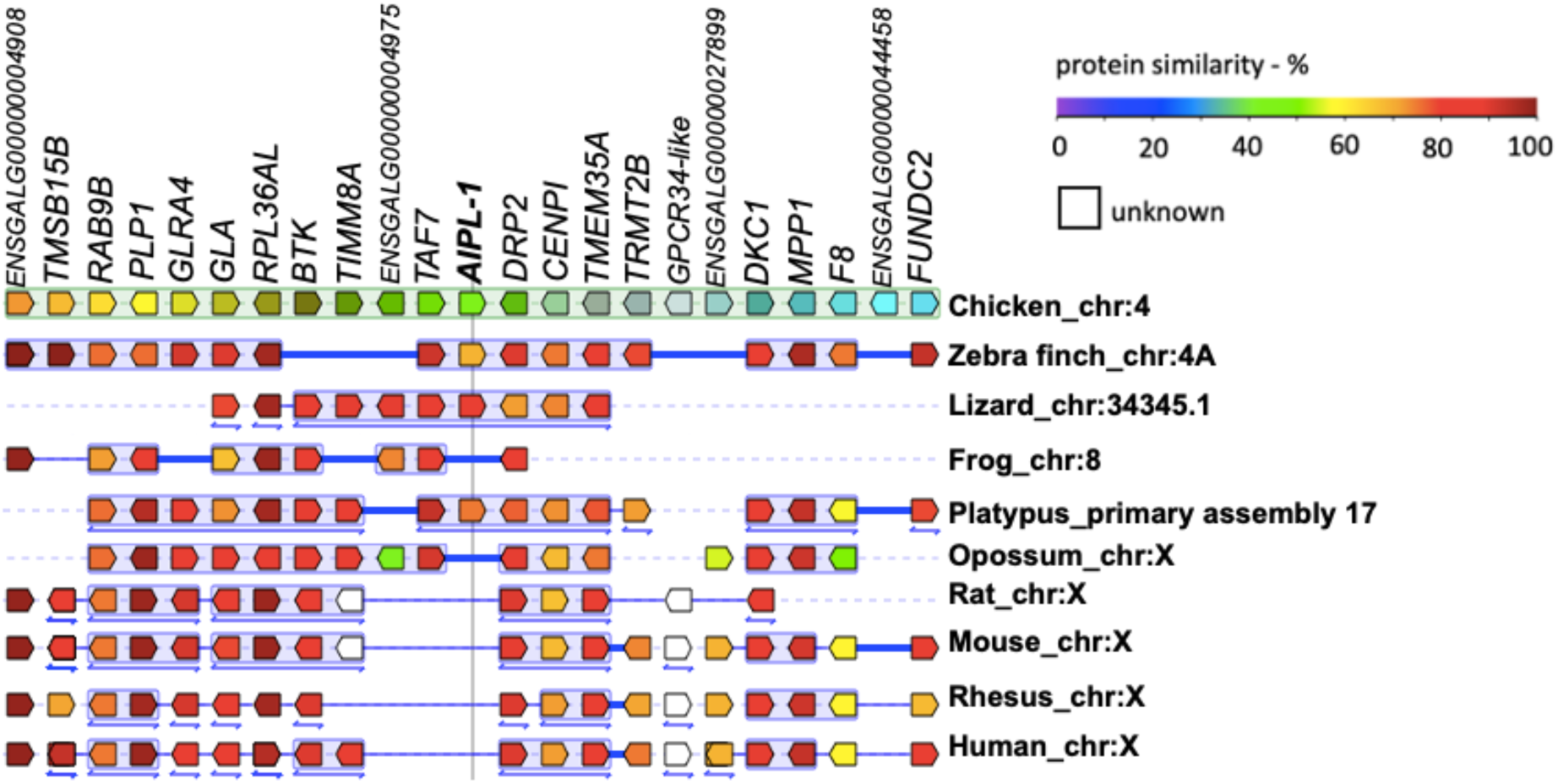
Synteny in the genomic neighborhood of chicken AIPL1 gene. The panel shows shared synteny with chicken AIPL1 (chromosome 4) as reference gene (vertical line). All these chromosomes fall on the same pre-1R paralogon described by (Lamb, 2021). The genes on either side of the reference gene are shown in correct order and orientation. Orthologs of the reference gene and its flanking genes are shown in the same color, below the reference chromosome, organized by species and chromosome. Consistent with the database, double-headed arrow under a block of genes indicates that the order of genes shown was flipped around. Thick blue line between genes is equivalent to a “gap” in the alignment of this species and thin blue line corresponds to a “break” of the alignment (two genes are linked in order, but at least one gene separates them in other species). Dotted line means no alignment. Synteny analysis was performed with Genomicus (Muffato et al., 2010) version 93.01 with the Alignview tool.

**Figure 9.**
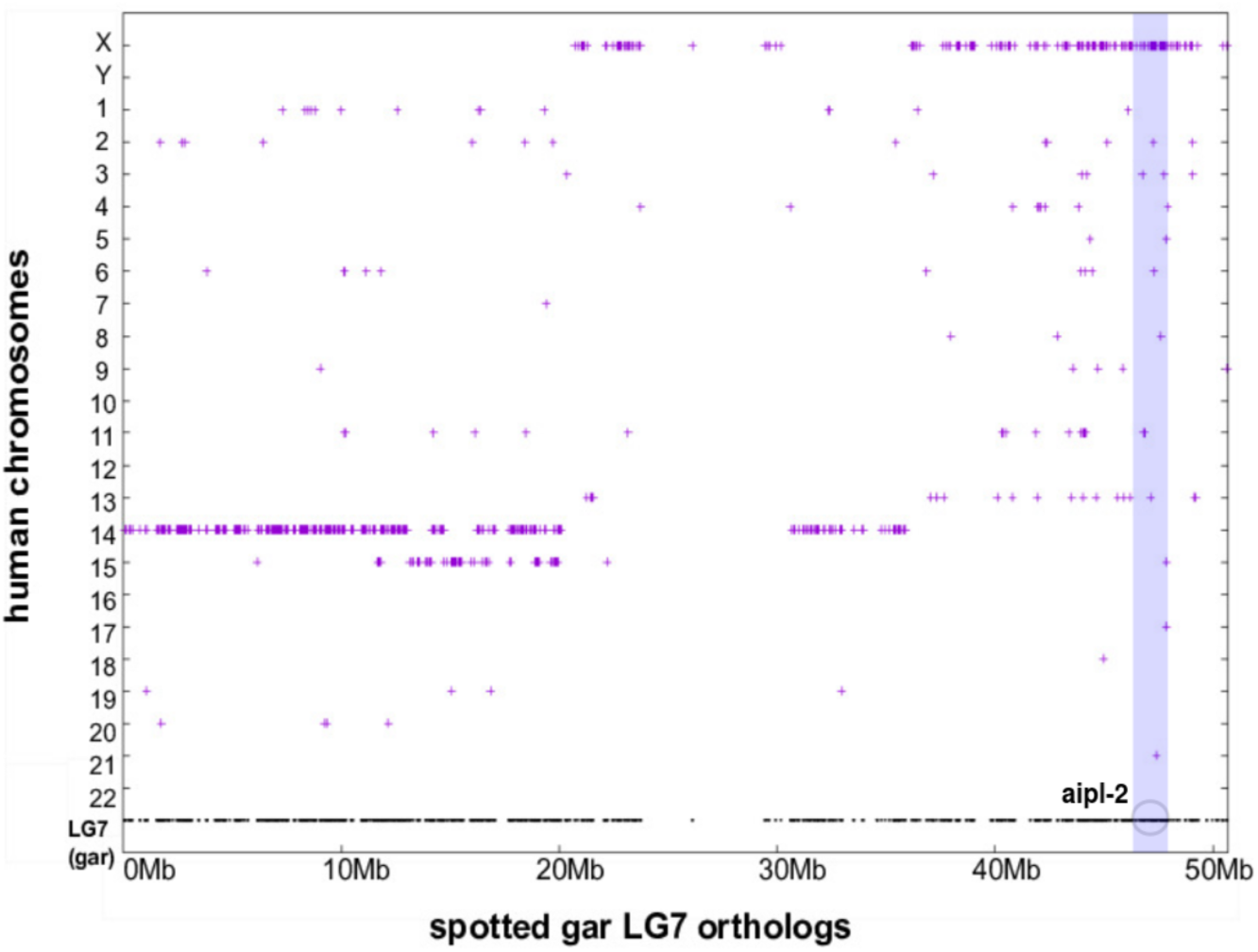
Orthology plot between spotted gar LG7 and human chromosomes. Dot plot showing the human orthologs of spotted gar LG7 genes. Human orthologs of LG7 are plotted vertically above the corresponding dot on their respective human chromosome. Location of spotted gar aipl-2 gene is highlighted in circle. LG7 shows strong orthology with HsaX and Hsa14 and weaker orthology with Hsa15. High degree of conservation along LG7 and HsaX supports that these occurred from a large-scale duplication event. Location of genes on LG 7 represent actual location on the chromosome, while human genes (orthologs to spotted gar LG7) on Hsa chromosomes do not necessarily represent their actual chromosomal location. Orthology dot plot created by Synteny Database.

## Discussion

AHR pathway plays an important role as sensor of environmental contaminants, closely inter-connected with multiple signaling pathways involved in early development and pathogenesis. This study illustrates evolutionary trajectory of AHR pathway genes and how multiple rounds of WGD have impacted its expansion. Our study shows strong indication that ancestral AHR pathway genes fall on pre-1R chromosome, eventually resulting on modern vertebrate paralogons in the current form as 2R-ohnologs (Figure 10). This points to possibility of these genes’ mechanistic relevance in disease as previous studies have described on 2R-ohnologue gene families’ involvement in disease progression through signaling pathways and transcription factors (Tinti *et al*., 2014; Singh *et al*., 2012; Huminiecki and Heldin, 2010). These findings will assist future studies on AHR related pathogenesis and in understanding human gene functions when using model organisms by supporting proper interpretation of comparative analysis.

**Figure 10.**
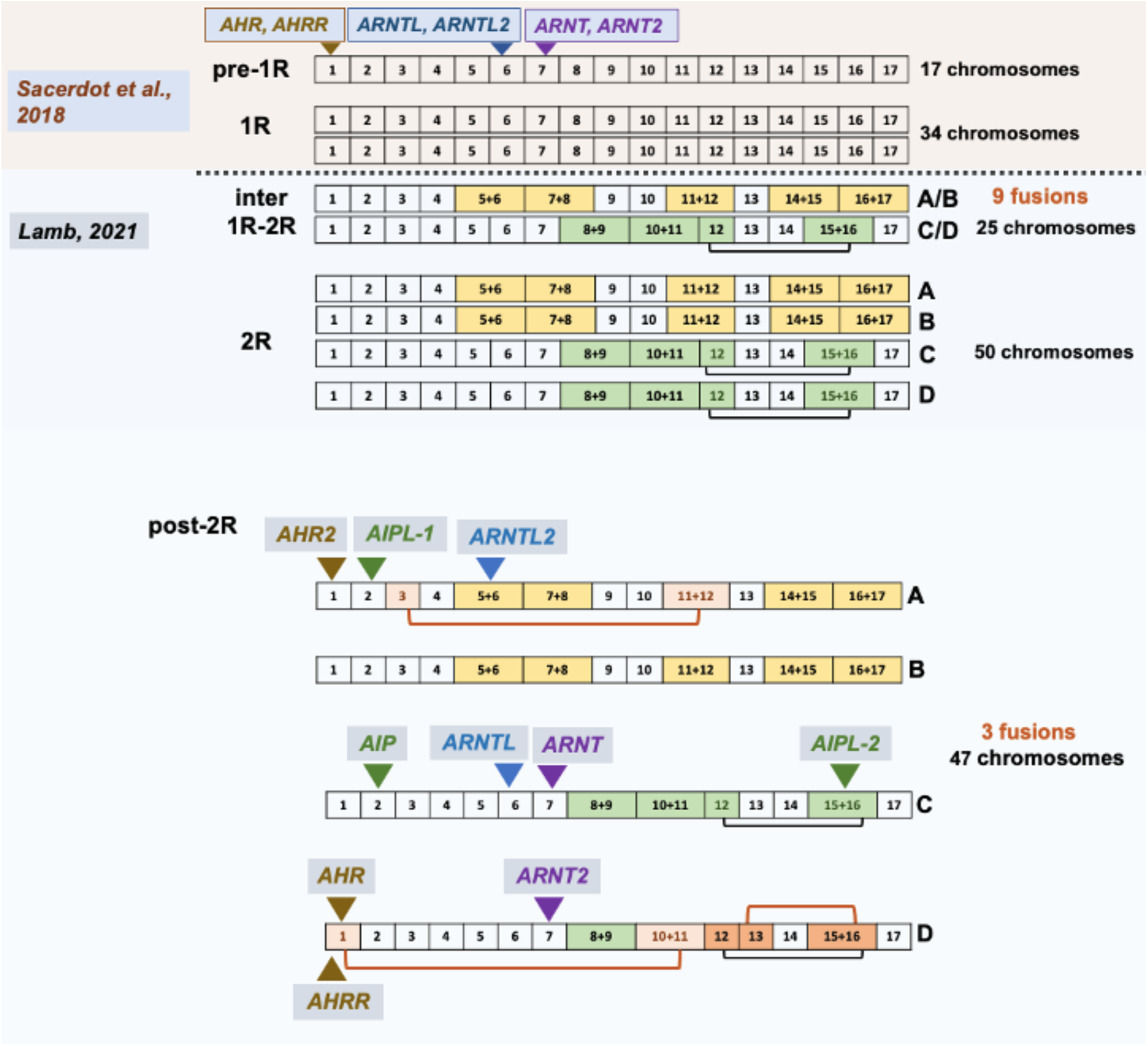
Summary of chromosomal location of AHR pathway genes. As summarized in Supplementary Table1, *Sacerdot et al*., *2018* map AHR pathway genes on the above pre-1R chromosomes. Horizontal dotted line indicates, two studies (*Sacerdot et al*. and *Lamb, 2021)* interpret differently below this. Post-2R is our analysis on AHR pathway genes’ chromosomal location mapped on to the paralogons inferred by *Lamb, 2021*. Orange line denotes chromosomal fusions post-2R before the radiation of bony-vertebrates.

Our analysis combining phylogenetic reconstruction followed by detailed synteny analysis revealed presence of AHR pathway genes in urochordates *Ciona*, predating two-rounds of WGD and traces of WGD on vertebrate chromosomes. Translational research using model organisms often suffers from poor cross-species extrapolation due to lack of proper evolutionary assignment of orthology. This study provides detailed analysis into the evolutionary mechanism of genes involved in AHR signaling, revealing WGD as the main contributor for its expansion. These findings are crucial for AHR research as recent years have seen expansion in this field and more importantly, our results point to its possible mechanistic relevance in disease caused by 2R-ohnologs through signaling pathways in multiple cancers and disease progression.

2R-ohnologs are particularly enriched in signaling molecules (as described by previous studies (Huminiecki and Heldin, 2010; Tinti *et al*., 2014)). The fact that an active pathway affecting multiple down-stream pathways such as the AHR signaling, expanded via WGD, resulting into a cascade of 2R-ohnologs, known to impact disease progression, has significant impact when studying these molecules and diseases related to this pathway.

Additionally, comparative analysis can assist in studies using model species to understand functional divergence following gene duplications (Conant and Wolfe, 2008) as previous studies on critical pathways have demonstrated (Cañestro *et al*., 2009).

Overall, our findings demonstrate main factors underlying molecular evolution of AHR pathway genes including its possible mechanistic relevance to disease due to 2R-ohnologs, which would be relevant to future studies in this field.

### Summary

This work illustrates evolutionary history of the genes involved in the AHR signal transduction pathway. AHR plays an important role in early development and response against environmental toxins. Sensitive homology searches with high quality sequence alignment followed by phylogenetic analysis suggests that this pathway was already present in the bilaterian ancestor in a primitive form and members have undergone duplications in the gnathostome lineage to give rise to the complete pathway. The reconciled phylogenies together with conserved synteny analysis reveal evolutionary traces of gene loss and teleost-specific duplications. Our analysis provides evidence of how AHR gene family evolution has been shaped by multiple WGDs and reconfirms that it was present at the base of vertebrate evolution. Further, it provides information essential for understanding functional diversity of multiple homologs of these gene families. This comparative approach will provide basis for understanding functional connectivity of human and model organisms that is crucial for translational research.

## Glossary

### XRE (xenobiotic response element)

An upstream regulatory sequence recognized by the transcription factors, particularly the aryl hydrocarbon receptor. These are conserved short DNA sequences in the promoter regions of many genes and the binding site for the receptor.

### Ohnolog

Gene duplicates originated by genome duplication.

## Acknowledgments

This research was supported by NC3Rs CRACK IT project DARTpaths (project number CRACKITDP-P1-2). Computational work was performed in the VSC (Flemish Supercomputer Center), funded by the Research Foundation - Flanders (FWO) and the Flemish Government.

